# Engineered human myogenic cells in hydrogels generate functional myofibers within dystrophic mouse muscle

**DOI:** 10.1101/2023.09.05.556335

**Authors:** Anna Kowala, Jinhong Meng, Olivier Pourquié, John Connelly, Jennifer E Morgan, Yung-Yao Lin

## Abstract

Transplantation of human myogenic progenitor cells (MPCs) is a promising therapeutic strategy for treating muscle-wasting diseases, e.g., Duchenne muscular dystrophy (DMD). To increase engraftment efficiency of donor stem cells, modulation of host muscles is required, significantly limiting their clinical translation. Here we develop a clinically relevant transplantation strategy synergising hydrogel-mediated delivery and engineered human MPCs generated from CRISPR-corrected DMD patient-derived pluripotent stem cells (PSCs). We demonstrate that donor-derived human myofibers produce full-length dystrophin at 4 weeks and 5 – 6 months (long-term) post transplantation in the unmodulated muscles of the dystrophin-deficient mouse model of DMD. Remarkably, human myofibers are innervated by mouse motor neurons forming neuromuscular junctions and supported by vascularisation after long-term engraftment in dystrophic mice. PAX7+ cells of human origin replenish the satellite cell niche. There was no evidence of tumorigenesis in mice engrafted with hydrogel-encapsulated human MPCs. Our results provide a proof-of-concept in developing hydrogel-based cell therapy for muscle-wasting diseases.

## Introduction

Skeletal muscle is responsible for the generation of all voluntary movements, such as walking and lifting objects. Skeletal muscle has a remarkable regenerative capacity mediated by resident muscle satellite cells ^1–3^. Muscular dystrophies are a group of genetic muscle-wasting conditions characterised by cycles of myofiber degeneration and regeneration with accumulation of fat and connective tissue. Duchenne muscular dystrophy (DMD), caused by dystrophin deficiency, is the most common form of muscular dystrophy in childhood, resulting in loss of ambulation, poor quality of life and premature death ^4^. Currently there is still no cure for any form of muscular dystrophies.

A potential approach for treating muscular dystrophies is cell-based therapy. A variety of stem/progenitor cells with myogenic potential have been used in previous studies to restore dystrophin expression in dystrophin-deficient mouse muscles, including satellite cells ^5,6^, myoblasts ^7,8^, pericytes ^9^ and human skeletal muscle-derived CD133+ (hCD133+) cells ^10,11^, as well as mesoangioblasts in dystrophin-deficient dogs ^12^. However, one of the major challenges in cell-based therapies for muscular dystrophies is obtaining sufficient number of stem/progenitor cells with myogenic potential for effective engraftment. This hurdle can now be addressed using human pluripotent stem cells (PSCs), embryonic stem cells (ESCs) or induced pluripotent stem cells (iPSCs), with transgene dependent or transgene-free differentiation protocols to generate an unlimited supply of myogenic progenitor cells (MPCs) for transplantation ^13,14^. Using a transcription factor PAX7-based fluorescence reporter, studies have shown that human PSC-derived MPCs are engraftable as they can not only give rise to dystrophin-positive myofibers, but also reconstitute the satellite cell compartment within the host mouse muscles ^15–17^.

Conventionally, donor cells with myogenic potential were transplanted into immunodeficient host animals through intramuscular cell injection. To increase engraftment efficiency, the host muscles were modulated prior to direct injection of donor cells using a range of regimens, including irradiation, cryoinjury, barium chloride (BaCl_2_) and myotoxins such as notexin and cardiotoxin. It was shown that deprivation of endogenous satellite cells in combination with preservation of host satellite cell niche significantly augmented donor mouse satellite cell engraftment ^18^. Apart from modulation of host muscles, it was shown that engraftment efficiency could also be affected by donor cell types and the recipient host animal strains that have different levels of immunodeficiency ^19^. Thus, both intrinsic properties of donor cells and local environment of host tissues are important factors in determining engraftment efficiency. Moreover, the microenvironment of dystrophic muscle may inhibit donor cell survival and differentiation as shown in the clinical trials ^20,21^. Despite progress demonstrated in previous proof-of-principle studies, delivery of donor cells via intramuscular injection is considered invasive and not practical in clinical settings as hundreds of injections would be required to cover large regions of skeletal muscle ^22,23^. In this regard, transplantation of donor cells via intramuscular injection significantly limits their clinical translation. Therefore, it is important to develop better transplantation methods to deliver human myogenic cells into host muscles for improving engraftment efficiency.

Tissue engineering approaches have been utilised to generate natural or synthetic biomaterial 3D scaffolds providing mechanical characteristics and signalling cues required for proliferation and differentiation of muscle stem/progenitor cells ^24–26^. To date, few studies have reported transplantation of human myogenic cell-laden hydrogels into host animals, including NOD scid gamma (NSG) mice ^27–29^, severe combined immunodeficient (SCID) mice ^30^, nude mice ^31^, and Rowett Nude (RNU) rats ^31,32^. In general, these studies reported the short-term (1 – 8 weeks) engraftment potential of the 3D scaffold systems with some promising results of innervation and vascularisation in the engrafted regions. However, it remains an open question whether the biomaterial 3D scaffolds can support engraftment of human myogenic cells in dystrophin-deficient dystrophic animal models by demonstrating that donor-derived human myofibers produce dystrophin. Furthermore, even though Rao et al. observed PAX7+ cells adjacent to myotubes within the implants at 2 – 3 weeks post transplantation ^28^, it remains to be demonstrated whether these PAX7+ cells were of donor or host origin or both, and whether PAX7+ cells of human origin could enter the satellite cell compartment in host animals. Finally, the long-term (up to 6 months) engraftment potential of human myogenic cell-laden hydrogels and their safety regarding tumorigenesis should also be investigated.

We previously generated two independent sources of human MPCs from two precisely CRISPR-corrected DMD patient-derived PSC lines (CORR-R3381X and CORR-K2957fs) and demonstrated the restoration of full-length dystrophin (Dp427) *in vitro* ^33,34^. In this study, we assess the *in vivo* regenerative potential of human CORR-R3381X or CORR-K2957fs MPCs compared with skeletal muscle-derived hCD133+ cells ^11^ by encapsulating them in fibrin/Matrigel hydrogels, followed by transplantation into dystrophin-deficient *mdx* nude mice. We show that engineered human myogenic cells contribute to muscle regeneration *in vivo* and express full-length dystrophin as demonstrated by quantification of human-specific antibodies against key markers. Importantly, we demonstrate innervation and vascularisation of human myofibers in the engrafted regions, PAX7+ cells of human origin repopulating the satellite cell niche and no evidence for tumorigenesis after long-term xenoengraftment (5 – 6 months). Together, our study provides a clinically relevant strategy synergising engineered human PSC-derived MPCs and hydrogel-mediated delivery for developing cell therapies to treat muscle-wasting conditions, such as DMD.

## Results

### Successful xenoengraftment by transplanting human cell-laden hydrogel constructs into dystrophin-deficient mice

To develop a clinically relevant transplantation protocol, we decided not to modulate host mouse muscles using irradiation, cryoinjury, BaCl_2_ or any myotoxin prior to donor human cell transplantation. In addition, we employed an established method to generate engineered 3D cell-laden constructs in polydimethylsiloxane (PDMS) molds ^24^ and investigated the regenerative potential of CRISPR-corrected human PSC-derived MPCs *in vivo*. Briefly, we encapsulated human myogenic cells in fibrin/Matrigel-based hydrogel cultured in growth medium (Day-2). The 3D cell-laden constructs remodelled within 2 days (Day 0). We then transplanted these 3D cell-laden constructs into dystrophin-deficient *mdx* nude mice (Day 3), followed by analysis at specific timepoints (Figure 1A). By placing hindlimbs in a position allowing access to the tibialis anterior (TA) muscles, we performed longitudinal skin and TA muscle incisions. In the meantime, the engineered 3D cell-laden constructs were cut with a 5 mm biopsy punch and removed from PDMS molds for transplantation. Next, a 3D cell-laden construct was embedded within the incision of the TA muscles and the skin incision was sutured (Figure 1B).

**Figure 1.**
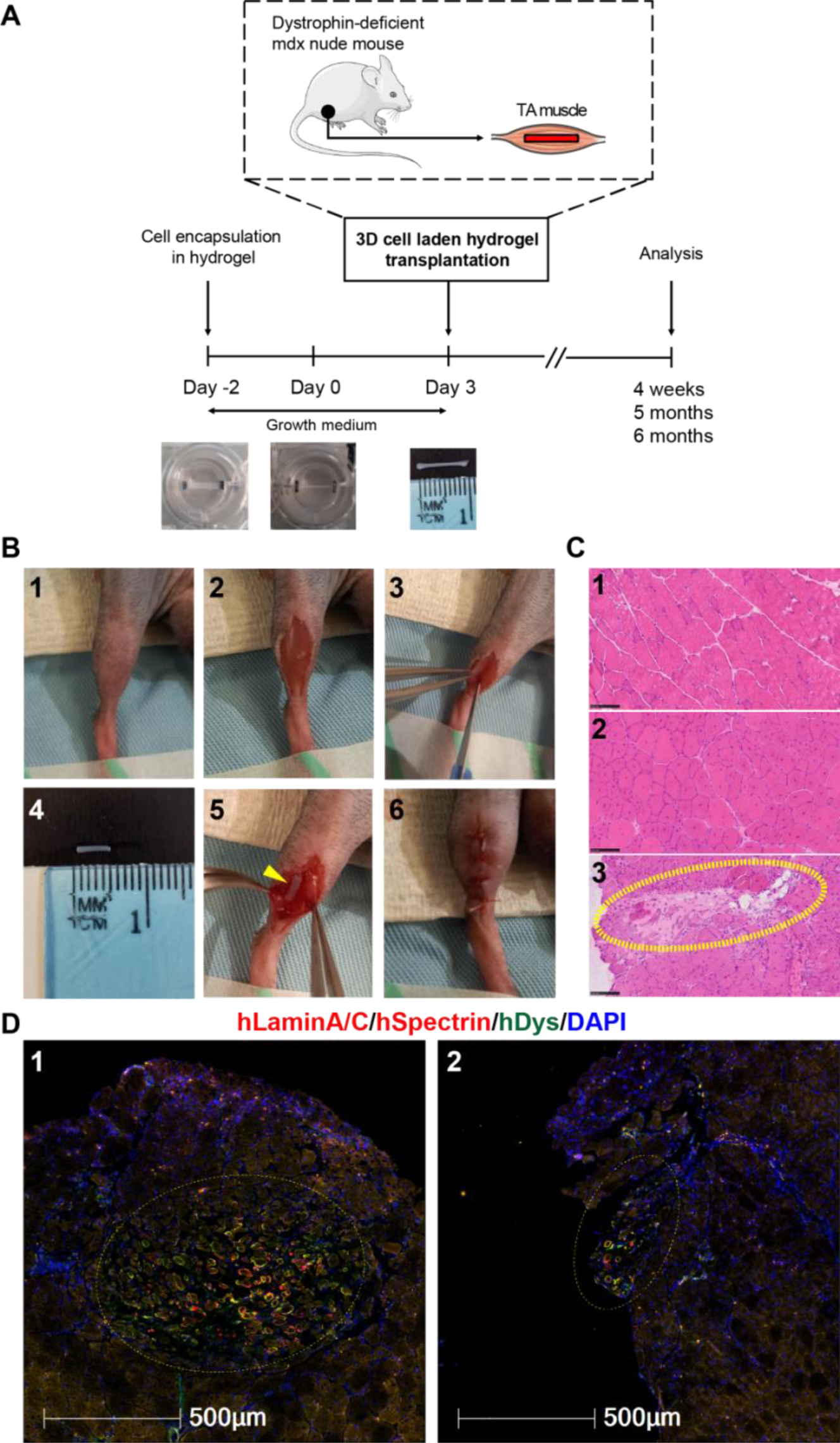
Experimental timeline and xenoengraftment of engineered human 3D cell-laden constructs in *mdx* nude mice. (A) Following encapsulation of human PSC-derived MPCs in hydrogel in PDMS molds, 3D cell-laden constructs were cultured for 5 days in growth medium (either Promocell or Megacell) and then transplanted into TA muscles of *mdx* nude mice, followed by analysis at 4 weeks, 5 months or 6 months post xenoengraftment. (B) Step-by-step transplantation procedure: 1) Positioning of the hindlimb; 2) Skin incision to reveal TA muscle; 3) TA muscle incision with a scalpel; 4) Prior to transplantation, hydrogel with encapsulated cells was cut with 5 mm-diameter biopsy punch and removed from its PDMS mold; 5) Placement of a 3D construct into TA muscle; and 6) Closure of the muscle incision, with a 3D construct inside, and suture of the skin. (C) Representative H&E staining of transverse cryosections of TA muscles. The hematoxylin (purplish blue) stains cell nuclei and eosin (pink) stains the extracellular matrix and cytoplasm. Panel 1, C57Bl/10 mouse (non-dystrophic control) with nuclei located at the periphery of myofibers; Panel 2, non-transplanted *mdx* nude (dystrophic control). Central nuclei within the myofibers are characteristic of *mdx* nude mouse muscles; Panel 3, at 4 weeks post transplantation, the transplanted hydrogel construct (dashed yellow circle) is visible within *mdx* nude TA muscle. Scale bars, 100 μm. (D) Representative images of hydrogel-based engraftment of human myogenic cells in TA muscle of *mdx* nude mice at 4 weeks post transplantation. Engrafted cells and myofibers of human origin in the middle of mouse TA muscle (dashed yellow circles, Panel 1) or at the edge of mouse TA muscle (dashed yellow circles, Panel 2). Transverse 10 μm sections were stained with antibodies against human lamin A/C and human spectrin (both red) and human dystrophin (green). Nuclei were counterstained with DAPI (blue). Scale bars, 500 μm.

As a control, we used healthy human skeletal muscle-derived hCD133+ cells (a subset of satellite cells), previously demonstrated capable of contributing to muscle regeneration and forming functional satellite cells after intramuscular transplantation in both immunodeficient Rag2-/γ chain-/C5- and *mdx* nude mice ^11,19^. For comparison with hCD133+ cells, we used two independent CRISPR-corrected human PSC-derived MPCs (CORR-R3381X and CORR-K2957fs). Prior to transplantation, cell-laden 3D constructs were cultured in different growth medium conditions for 5 days. At 4 weeks post transplantation, hematoxylin and eosin (H&E) staining showed the presence of hydrogel implants within TA muscle of *mdx* nude mice (Figure 1C). Among 56 analysed TA muscles, immunostaining of muscle sections with human-specific antibodies (hLaminA/C, hSpectrin and hDystrophin) revealed myofibers of human origin either in the middle (64.29%) or at the edge (32.14%) of the host TA muscle (Figure 1D). Only 3.57% of analysed TA muscles contained no myofibers of human origin. Importantly, transverse sections of host muscles contained not only hLaminA/C+ nuclei (of human origin) but also hSpectrin+ and hDystrophin+ myofibers (Figure 1D), demonstrating *in vivo* skeletal muscle regeneration from engineered human myogenic cells. Together, these results indicate successful xenoengraftment of human myogenic cells in dystrophin-deficient *mdx* nude mice by hydrogel-mediated delivery.

### CORR-R3381X and CORR-K2957fs MPCs contribute to muscle regeneration *in vivo*

similar to skeletal muscle-derived hCD133+ cells Next, we sought to compare the efficiency of *in vivo* muscle regeneration between different experimental conditions by quantifying myofibers of human origin. Notably, it was shown that newly regenerated myofibers in *mdx* nude, *NOD/Rag1(null)mdx(5cv)*, or *NOD/LtSz-scid IL2Rγ(null)* mice were recognised by anti-human spectrin antibody, which might be due to expression of utrophin in regenerating myofibers ^35^. This might lead to detection of false-positive donor myofibers. Moreover, after transplantation into host muscles, myogenic cells might not only fuse with each other forming myofibers of donor origin, but also fuse with host myogenic cells or host myofibers, resulting in mosaic myofibers ^36^. Therefore, spectrin and dystrophin immunostaining might show a mosaic pattern ^36,37^. To interpret data cautiously, we decided to quantify myofibers of human origin as hSpectrin+ myofibers containing hLaminA/C+ nuclei, and any hDystrophin+ myofibers. In addition, hLaminA/C+ nuclei that were not within a myofiber containing hSpectrin or hDystrophin might have been either satellite cells, or undifferentiated cells of human origin.

In total we compared 5 experimental conditions, in which CORR-R3381X and CORR-K2957fs cell-laden constructs were cultured in either Promocell or Megacell growth medium for 5 days after encapsulation, whereas hCD133+ cell-laden constructs were cultured in Megacell growth medium. In agreement with previous studies ^11,19^, we showed that all hCD133+ cell-laden constructs contributed to muscle regeneration *in vivo* at 4 weeks post transplantation (Figure 2A; Supplementary Table S1). Nonetheless, it should be noted that the number of human myofibers vary widely in the host mice, ranging from 2 to 277 human myofibers in host TA muscles (Figure 2B; Supplementary Table S1), which was consistent with the transplantation efficiency of the same type of cells (without hydrogel encapsulation) into irradiated and cryoinjured *mdx* nude mouse muscle ^11,19^. In contrast, CORR-R3381X and CORR-K2957fs cell-laden constructs cultured in Megacell medium generated up to 7 and 10 human myofibers, respectively, in one host TA muscle (Figure 2B; Supplementary Table S1). However, we found that both CORR-R3381X and CORR-K2957fs cell-laden constructs cultured in Promocell medium gave rise to higher number of donor-derived human myofibers than those cultured in Megacell medium (Figure 2B). Among three independent sets of transplantations, CORR-R3381X constructs generated up to 177 human myofibers in one host TA muscle, whereas CORR-K2957fs constructs could produce up to 60 human myofibers in one host TA muscle (Supplementary Table S1). Among the independent experiments, the greatest number of myofibers of human origin per host TA muscle (median [25th – 75th percentile]) from hCD133+, CORR-R3381X and CORR-K2957fs cell-laden constructs were 78 [33.5 – 160.8], 69 [24.75 – 111], and 24 [11.5 – 42.5], respectively (Figure 2B; Supplementary Table S1). Next, we pooled three sets of data together to investigate the percentages of undifferentiated cells (hLaminA/C+ only) and myofibers of human origin generated by CORR-R3381X or CORR-K2957fs constructs in Promocell medium, and hCD133+ constructs in Megacell medium. In general, CORR-R3381X and hCD133+ constructs showed very similar percentages of undifferentiated cells (∼11.72 – 12.9%) and human myofibers (∼87.1 – 88.28%), whereas CORR-K2957fs constructs appeared a higher percentage of undifferentiated cells (∼21.69%) and lower percentage of human myofibers (∼78.31%) (Figure 2C).

**Figure 2.**
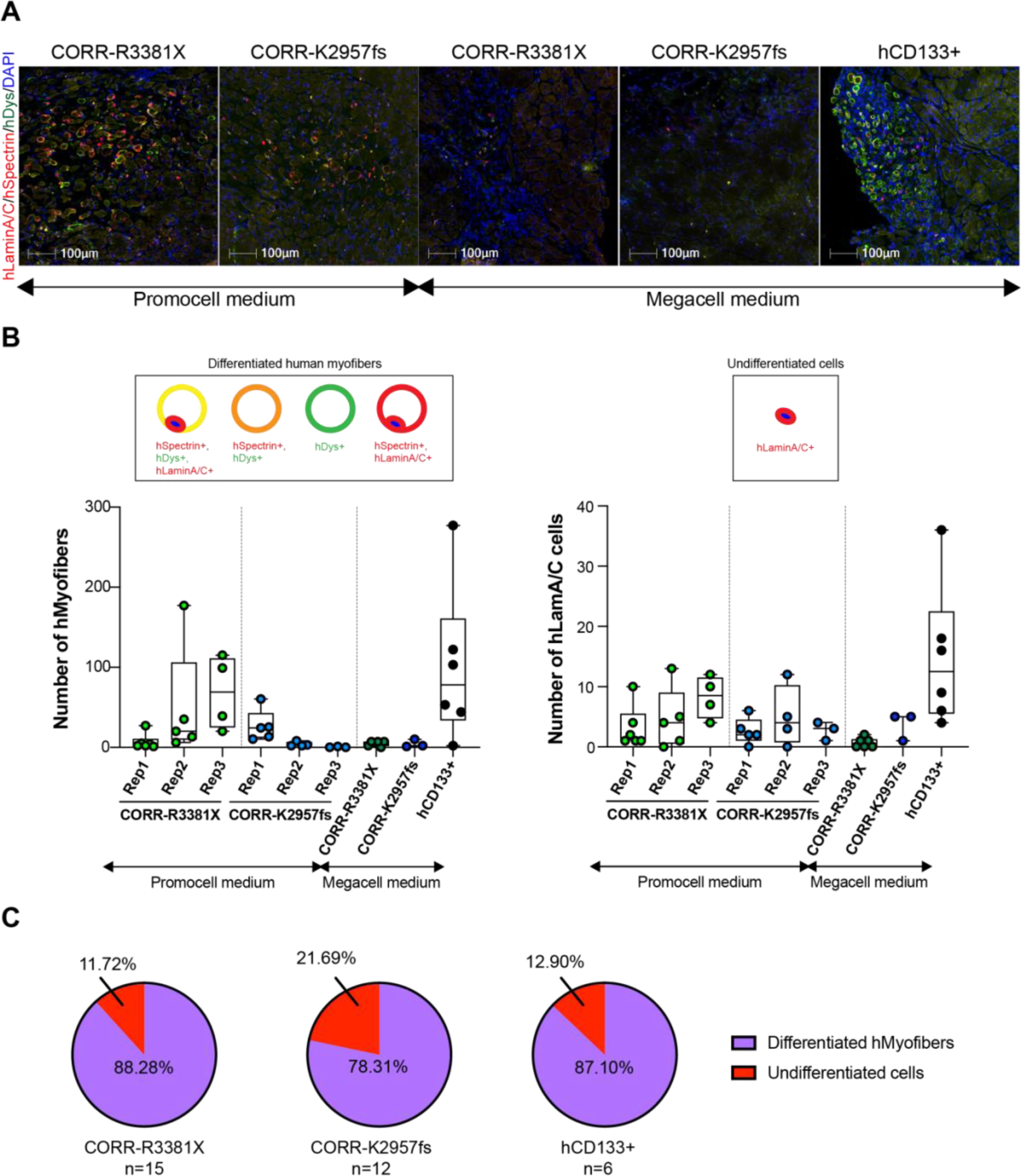
Engrafted cells and myofibers of human origin in *mdx* nude mice at 4-week post transplantation. (A) Representative transverse cryosections of *mdx* nude mouse TA muscle transplanted with 3D constructs of CORR-R3381X MPCs in Promocell growth medium; CORR-K2957fs MPCs in Promocell growth medium; CORR-R3381X MPCs in Megacell growth medium; CORR-K2957fs MPCs in Megacell growth medium; hCD133+ cells in Megacell growth medium. At 4 weeks post transplantation, transverse 10 μm sections were stained with antibodies against human lamin A/C and human spectrin (both red) and human dystrophin (green). Nuclei were counterstained with DAPI (blue). Scale bars, 100 μm. (B) Quantification of myofibers and undifferentiated cells of human origin in each experimental condition or independent replicate (Rep 1, 2 and 3). Schematics represent immunostaining patterns considered as myofibers of human origin (hMyofibers) or undifferentiated cells of human origin (hLamin A/C+ only). (C) Pie charts show percentages of engrafted human myogenic cells in *mdx* nude mice as undifferentiated cells and differentiated human myofibers. CORR-R3381X MPCs in Promocell growth medium (15 TA muscles); CORR-K2957fs MPCs in Promocell growth medium (12 TA muscles); hCD133+ cells in Megacell growth medium (6 TA muscles).

Taken together, even though the engraftment efficiencies vary widely between cell-laden constructs within the same experiment, as well as between constructs in different sets of experiments, our results demonstrated that CORR-R3381X and CORR-K2957fs MPCs can contribute to muscle regeneration *in vivo*, like skeletal muscle-derived hCD133+ cells. The fact that cell-laden constructs maintained in Promocell medium gave markedly higher engraftment efficiency than those maintained in Megacell medium suggest that engraftment efficiency could be further improved by modulating human myogenic cells with appropriate growth medium prior to transplantation.

### CORR-R3381X and CORR-K2957fs MPCs gave rise to similar distribution of mosaic patterns of donor-derived human myofibers as hCD133+ cells

We subsequently investigated whether the number of donor-derived human myofibers are similar between CORR-R3381X, CORR-K2957fs and hCD133+ cell-laden constructs. To do this, we quantified the number of donor-derived human myofibers in 4 categories, including i) hDystrophin+ only; ii) hSpectrin+ with hDystrophin+ and hLaminA/C+; iii) hSpectrin+ with hDystrophin+; and iv) hSpectrin+ with hLaminA/C+ (Figure 3A; Supplementary Table S1). Regardless of the myogenic cell sources and engraftment efficiencies, we found that >99.29% donor-derived human myofibers in dystrophin-deficient *mdx* nude mouse TA muscles express human dystrophin (Figure 3B). Moreover, CORR-R3381X and CORR-K2957fs constructs gave rise to very similar percentage distribution of human-specific markers in myofibers in post transplantation TA muscle sections with the donor-derived human myofibers expressing hSpectrin+ with hDystrophin+ (∼62.12 – 73.65%), hSpectrin+ with hDystrophin+ and hLaminA/C+ (∼25 – 32.39%), or hDystrophin+ only (∼1.35 – 4.78%) (Figure 3B). These results suggest that CRISPR-corrected human PSC-derived MPCs may contribute to muscle regeneration after xenotransplantation in a similar way to other muscle precursor cells of human origin, such as hCD133+ cells.

**Figure 3.**
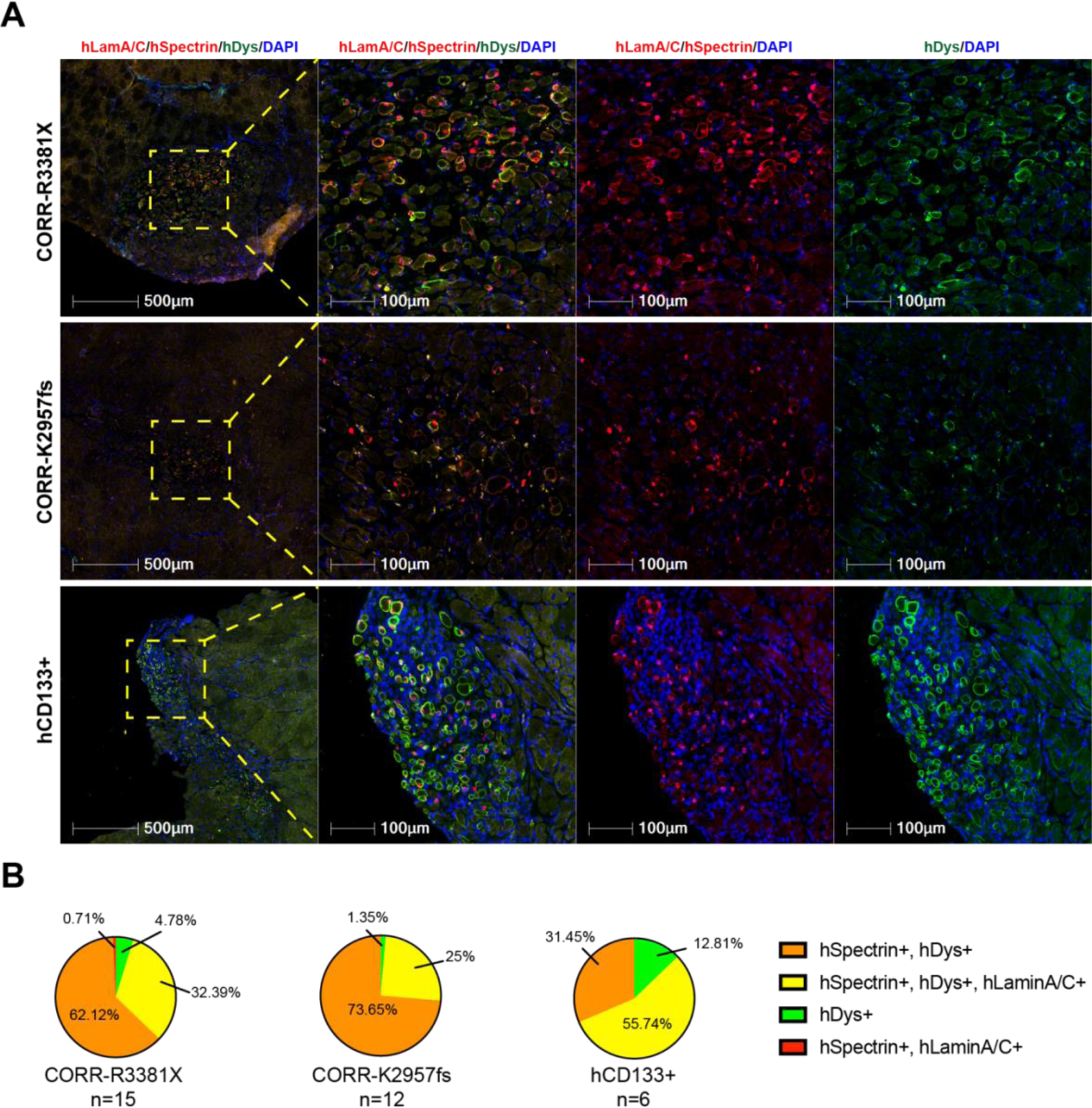
Distribution patterns of donor-derived human myofibers in *mdx* nude mice at 4 weeks post transplantation. (A) Representative transverse cryosections of donor-derived human myofibers derived from cell-laden constructs of CORR-R3381X MPCs or CORR-K2957fs MPCs in Promocell growth medium, and hCD133+ cells in Megacell growth medium. Human lamin A/C and human spectrin (both red) and human dystrophin (green). Nuclei were counterstained with DAPI (blue). Scale bars, 100 μm. (B) Pie charts show percentages of distribution patterns of donor-derived human myofibers in *mdx* nude mice. CORR-R3381X MPCs in Promocell growth medium (15 TA muscles); CORR-K2957fs MPCs in Promocell growth medium (12 TA muscles); hCD133+ cells in Megacell growth medium (6 TA muscles).

### Long-term engraftment of human myofibers with replenished satellite cell niche supported by innervation and vascularisation in *mdx* nude mice

Next, we sought to test whether 3D cell-laden constructs might support long-term engraftment of human myogenic cells and whether there is any concern of tumorigenesis. Given that CORR-R3381X cell-laden constructs gave better engraftment efficiency than CORR-K2957fs constructs at 4 weeks post transplantation, we decided to focus on transplantation of CORR-R3381X constructs cultured in Promocell medium, followed by analysis at either 5 (n=1) or 6 (n=5) months. In total, we transplanted 6 cell-laden constructs into 6 TA muscles of *mdx* nude mice. Encouragingly, donor-derived human myofibers were detected in one TA muscle that was analysed at 5 months post transplantation (98 human myofibers) and two TA muscles at 6 months post transplantation (up to 59 human myofibers), as evidenced by hDystrophin+ staining (Figure 4; Supplementary Table S2). Importantly, there is no evidence for tumorigenesis in any mouse transplanted with hydrogel-encapsulated human MPCs after long-term engraftment.

**Figure 4.**
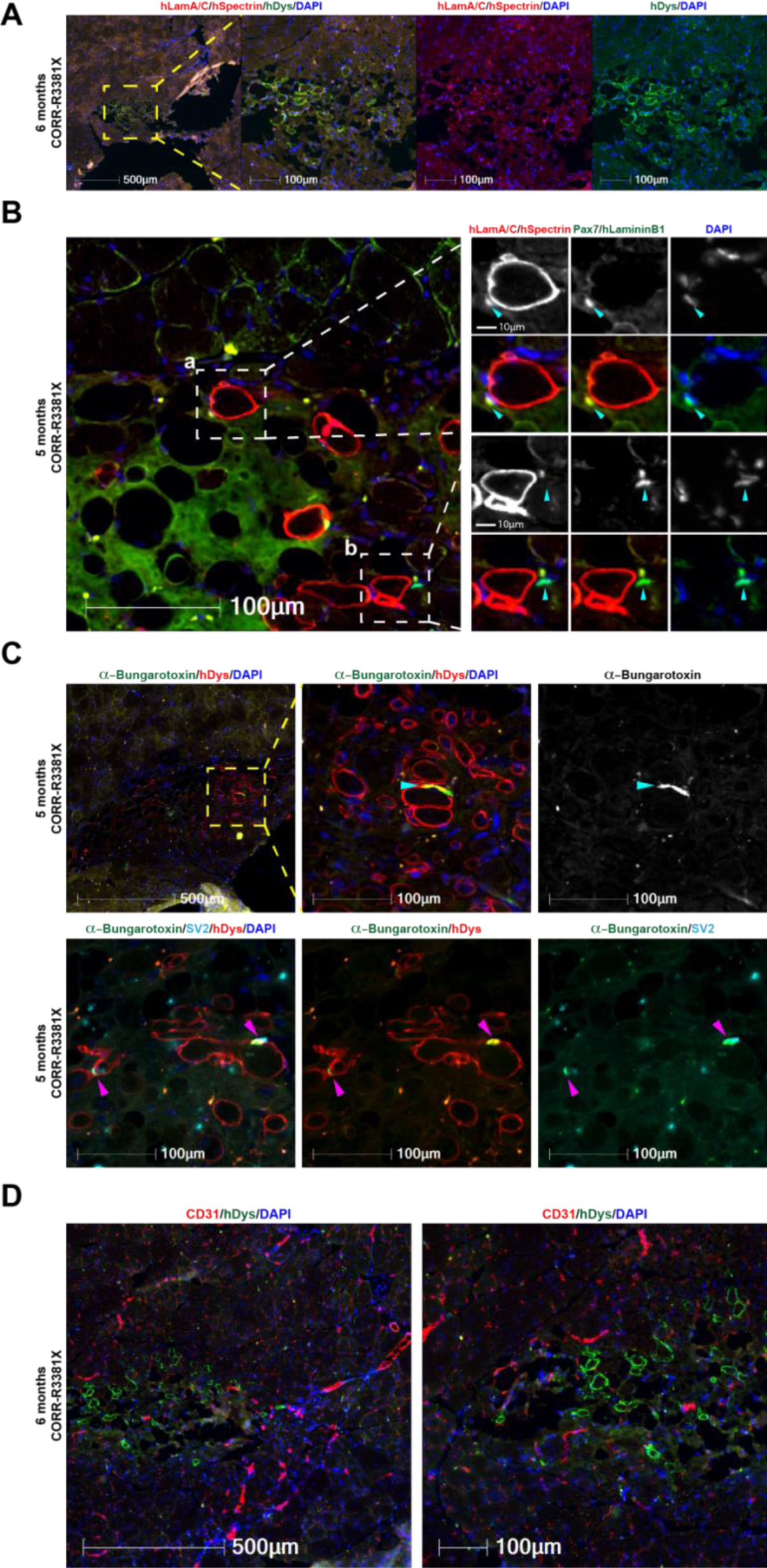
Long-term engraftment of human myofibers with repopulation of satellite cell niche supported by innervation and vascularisation in *mdx* nude mice. (A) A representative transverse cryosection of *mdx* nude mouse TA muscle transplanted with a 3D construct of CORR-R3381X MPCs in Promocell growth medium (6 months post transplantation). Human lamin A/C and human spectrin (both red); human dystrophin (green). Nuclei labelled with DAPI (blue). (B) Detection of human (a) and mouse (b) PAX7+ cells (arrowheads) in the satellite cell compartment at 5 months post transplantation. (a) A PAX7+ cell (green) of human origin labelled by human laminA/C (red) and DAPI (blue) adjacent to a human myofiber (hSpectrin, red) beneath the basal lamina (hLaminin β1, green). (b) A PAX7+ cell (green) of mouse origin with DAPI (blue) adjacent to a human myofiber (hSpectrin, red) within the basal lamina (hLaminin β1, green). (C) Innervation of donor-derived human myofibers (hDystrophin, red) in *mdx* nude mice at 5 months post transplantation. The formation of NMJs (arrowheads) is demonstrated by co-localisation of post-synaptic marker AChR (labelled by α-Bungarotoxin, green) and pre-synaptic marker SV2 (cyan). Nuclei labelled with DAPI (blue). (D) Vascularisation in the engrafted regions at 6 months post transplantation as demonstrated by detection of blood vessels (CD31, red) adjacent to human myofibers (hDystrophin, green). Nuclei (DAPI, blue). Scale bars, 10 – 500 μm as indicated in each panel.

By examining transverse cryosections stained with antibodies against hLaminin B1, hSpectrin and hLamin A/C, as well as the canonical satellite cell marker PAX7, we identified hLamin A/C and Pax7 double-positive cells of human origin residing in the satellite cell niche (underneath the basal lamina of myofibers) in the TA host muscle (Figure 4B, Panel a). In addition, we could also identify cells that were hLamin A/C-negative and Pax7-positive close to the engrafted region in mouse TA muscle, indicating these were satellite cells of mouse origin (Figure 4B, Panel b). Together, these results suggest that transplantation of 3D cell-laden hydrogel constructs not only contribute to muscle regeneration *in vivo*, but also replenish the satellite cell compartment in the host muscle.

Given the long-term persistence of human myofibers in *mdx* nude mice, we hypothesise that the human myofibers might be supported by innervation and vascularisation in the host muscle tissue. By using fluorescence-conjugated α-Bungarotoxin to label the post-synaptic acetylcholine receptor (AChR) and antibodies against the pre-synaptic marker synaptic vesicle glycoprotein 2A (SV2), we detected not only AChR clustering on hDystrophin+ myofibers (Figure 4C), but also the co-localisation of AChR and SV2 indicating the formation of neuromuscular junctions (NMJs) between mouse motor neuron and human myofibers (Figure 4C). Furthermore, using CD31 antibodies that recognise endothelial cells, we identified blood vessels spreading across the engrafted region of hDystrophin+ myofibers (Figure 4D), supporting our hypothesis. Taken together, these results suggest that transplantation of 3D cell-laden hydrogel constructs supports long-term engraftment of human myofibers that are innervated and vascularised, together with a replenished satellite cell niche, in *mdx* nude mice.

## Discussion

In this study, we have established a clinically relevant transplantation strategy that uses hydrogel-mediated deliveries of engineered human myogenic cells without modulation of the host muscles to achieve xenoengraftment in dystrophin-deficient *mdx* nude mice. We have shown that two independent lines of CRISPR-corrected human PSC-derived MPCs and skeletal muscle-derived hCD133+ cells contribute to *in vivo* muscle regeneration with expression of full-length dystrophin. For the first time, we have demonstrated innervation and vascularisation of donor-derived human myofibers within *mdx* nude mouse muscles at 5 – 6 months post transplantation. Moreover, engrafted human myogenic cells gave rise to PAX7+ cells to replenish satellite cell niche. Importantly, there was no evidence for tumorigenesis in transplanted mice. Together, our results suggest that human PSC-derived MPCs are safe and stable for long-term engraftment. Transplantation of autologous MPCs rather than allogenic donor-derived MPCs reduces risk of the immunological response leading to rejection and reduce necessity for lifelong immunosuppression. Our findings suggest that it is possible to use CRISPR-corrected PSCs to generate MPCs to develop hydrogel-based cell therapy and our strategy can be applied to other types of muscular dystrophy ^38^. But our study also shows a wide variation of engraftment efficiencies between sources of myogenic cells, between individual host mice, and between transplantation experiments. Therefore, there is still room for further improvement to reduce the variability and increase the engraftment efficiency.

Several factors may affect the variability of engraftment efficiency. For instance, quantification of myofibers of human origin could be affected by technical aspects related to the experimental and analysis methods. In 10 µm sections of analysed host *mdx* nude TA muscles, we could not always detect nuclei in all the donor-derived human myofibers. In addition, we used a stringent quantification method, which excluded hSpectrin+ only myofibers as human origin if they did not have hLaminA/C+ nuclei, nor hDystrophin+ immunofluorescence. Thus, our study is likely to underestimate myofibers of human origin. Another pertinent factor is that transplanted hydrogel constructs may have been squeezed either to the edge or outside the grafted muscle, which might affect the engraftment efficiency. After immunofluorescence analysis with human-specific antibodies, we observed 32.14% of transplants at the edge of the host muscle, which indicate sub-optimal hydrogel placement, resulting in technical issues with cryosectioning and further analysis. For example, we observed that during cryosectioning the outer part of the muscle section occasionally detached from the rest, which could also have a negative impact on our quantification. Notably, even though nude mice are T-cell deficient, they maintain B cell activity and high NK cell function ^39,40^. In this respect, low number of donor-derived human myofibers in *mdx* nude mice might be because xenografts suffered some immunological rejection in host mice which are not completely immunodeficient. Moreover, previous data suggested that genetic background has an influence on *mdx* mouse muscle regeneration ^41^. Therefore, differences of genetic backgrounds between our *mdx* nude mice and other groups’ dystrophin-deficient immunodeficient host mice might also affect the number of myofibers of human origin in different studies.

Apart from host animal strains ^19^, it was shown that modulation methods of host muscle can also affect the engraftment efficiency, e.g., prior to transplantation of donor cells, host muscles can be irradiated, cryoinjured or both ^19^. Irradiation has been used to recapitulate the DMD pathology in *mdx* mice, in which satellite cell exhaustion hinders muscle regeneration ^42^. In addition, irradiation spares myofibers and preserves the muscle stem cell niche and has an influence on regenerative potential of transplanted cells ^43^. Indeed, previous studies showed that significantly more donor-derived dystrophin+ myofibers were present in irradiated than in non-irradiated host muscles after immortalised mouse myoblasts ^44^ or mouse satellite cells ^5,43,45^ were transplanted into *mdx* nude mice, but host muscle irradiation did not augment muscle regeneration derived from human muscle precursor cells ^19^. Notably, childhood cancer survivors have shown that radiation exposure in early life stage can cause radiation-induced fibrosis affecting 80% of patients ^46^, leading to muscle atrophy, impaired mobility and weakness ^47^. Thus, it is desirable to develop new clinically safe approaches boosting donor cell engraftment in future cell therapies for DMD patients.

Previous studies have shown regenerative potential of mouse ^6,43^ or human ^48^ satellite cells after transplantation into host mouse muscles. These satellite cells demonstrated engraftment and generation of myofibers of donor origin within the host muscles. It has been also shown that intramuscular transplantation of human postnatal myoblasts, skeletal muscle-derived hCD133+ cells, or human PSC-derived MPCs into immunodeficient mice can give rise to functional satellite cells of donor origin ^11,16,49^. We hypothesise that PAX7+ cells of human origin post transplantation in our study may be functionally equivalent to satellite cells. However, this will require further investigation to demonstrate whether these PAX7+ cells of human origin in the satellite cell niche are able to contribute to host muscle regeneration after re-injury.

Our study has some limitations. Firstly, as we used Matrigel mixed with fibrin for cell encapsulation to generate hydrogel constructs. Since Matrigel is a solubilised basement membrane preparation derived from the Engelbreth-Holm-Swarm mouse sarcoma, many inherent issues are associated with Matrigel, such as lot-to-lot variation of extracellular matrix composition, potential pathogen transmission and risks for immunogenicity in humans. Thus, these issues prohibit its clinical use and highlight the need to develop xeno-free natural hydrogels (e.g., protein polymers; decellularised extracellular matrix) or synthetic biomaterials (e.g., polyethylene glycol macromer) ^25,26^ suitable for hydrogel-mediated delivery of human MPCs with improved engraftment efficiency. Secondly, in our study, the percentage of donor-derived human myofibers per mouse TA muscle (∼2000 myofibers) is lower than 20% in any set of transplantations. Although it seems only modest levels of dystrophin are required for functional benefit ^50–55^, to be fully protective, dystrophin at low levels must spread all along the myofiber ^56^ and present in most of the fibres within a muscle. However, the contribution of donor cell-derived dystrophin to host mouse muscle is segmental ^36,37,57^ and the spreading of dystrophin is only a few hundred microns from the nucleus producing it ^36^. For these reasons, we did not perform experiments to assess functional improvements in *mdx* nude TA muscles transplanted with CORR-R3381X or CORR-K2957fs cell-laden constructs. Nevertheless, when future studies achieve 10 – 20% of dystrophin present in the majority of myofibers within the treated muscle, it will be of interest to assess whether xenoengraftment is able to improve muscle function of dystrophin-deficient host mice.

In conclusion, we have shown that hydrogel-mediated delivery is suitable for long-term engraftment of CRISPR-corrected human PSC-derived MPCs in unmodulated *mdx* nude mouse muscles, enabling full-length dystrophin expression, innervation and vascularisation in the engrafted region, and repopulating the satellite cell niche. In this regard, our study therefore presents a clinically relevant transplantation strategy. Advances in xeno-free natural or synthetic biomaterials that can replace Matrigel-based hydrogels and support regenerative potential of human MPCs will pave ways for clinical trials of hydrogel-mediated cell therapy. Future improvement in engraftment efficiency with evidence of improved muscle function may lead to therapeutic application for muscular dystrophies.

## Materials and Methods

### Ethics

Under appropriate ethical approvals by Hammersmith and Queen Charlotte’s and Chelsea Hospital (REC reference 06/Q0406/33) and by National Research Ethics Service Committee London-Stanmore (REC reference 13/LO/1826; IRAS project ID: 141100), informed consent was obtained prior to the use of human derived cells.

### Human myogenic cells

Healthy human skeletal muscle-derived CD133+ cells (hCD133+) were described ^11^ and obtained from Medical Research Council Centre for Neuromuscular Diseases Biobank (ID: 8206). CORR-R3381X and CORR-K2957fs human MPCs were generated using a transgene-free myogenic differentiation protocol ^58^ from two precisely CRISPR-corrected DMD patient-derived PSC lines, respectively ^33,34,59^. hCD133+ cells were maintained in Megacell Skeletal Muscle Growth Medium (For details, see Supplementary Table S3), whereas CORR-R3381X and CORR-K2957fs MPCs were maintained in Promocell Skeletal Muscle Growth Medium (Promocell, C-23060).

### *Mdx^nu/nu^* (*mdx* nude) mouse

Host mice aged 5 to 8 weeks were used for this study. A well-established immunodeficient *mdx* nude mouse model ^8,44^, which harbours a nonsense mutation at the exon 23 of the dystrophin gene, lacking T cells and partial B cell deficiency, was used as host. All animal experiments were conducted at the Biological Services Unit, University College London Great Ormond Street Institute of Child Health, in accordance with the Animals Act 1986, approved by the University College London Animal Welfare Ethical Review Body. Experiments were performed under Home Office licence number: PP2611161.

### Cell encapsulation in fibrin/Matrigel hydrogel in PDMS molds

Upon cell encapsulation, CORR-R3381X and CORR-K2957fs human MPCs were expanded in Promocell medium, whereas hCD133+ cells were expanded in Megacell Medium for around 2 weeks with passage number P7 – P10. The 24-well format PDMS molds were fabricated as described, followed by addition of Velcro pieces at both ends of dumbbell-shaped depression in the molds ^24^. Fibrin/Matrigel hydrogel mixture was prepared according to the recipe (Supplementary Table S3). 3D cell-laden constructs were prepared by encapsulating 10 x 10^6^ cells per ml hydrogel mixture, followed by adding thrombin (Sigma, T6634; 0.2 unit per mg of fibrinogen) prior to evenly seeding cell/hydrogel suspension in PDMS molds, resulting in 0.5 x 10^6^ cells/construct. 3D cell-laden constructs were then cultured in skeletal muscle growth medium. 5 days after cell encapsulation, 3D cell-laden constructs were cut with biopsy punch (Stiefel, D5245) to 5 mm in length prior to transplantation into *mdx* nude mice.

### *In vivo* transplantation procedure of 3D cell-laden constructs

Surgery was performed under sterile conditions. Immunocompromised *mdx* nude mice (between 5 and 8 weeks old) were anaesthetised with isoflurane and injected subcutaneously with Metacam analgesic (5 mg/kg). The mice were kept warm during surgery that involved a longitudinal skin incision to expose the tibialis anterior (TA) muscle. Afterwards, a disposable scalpel (Swann-Morton, 0503, size 11) was used to cut the TA muscle longitudinally and a 5-mm long 3D cell-laden construct was placed inside the incision. The incision was closed carefully, making sure the construct remained within the TA muscle, and the skin was sutured (polyglactin 910, 6-0 coated, P-1, Vicryl Rapide, W9913). All mice recovered within half an hour post transplantation without any post-operative complications.

### TA muscle tissue processing

Mouse TA muscles were dissected at 4 weeks, 5 months or 6 months post transplantation, and embedded in 6% gum tragacanth (Sigma G-112) on cork disks, before being frozen in pre-chilled isopentane in liquid nitrogen. Serial 10 µm transverse cryosections were cut throughout the muscle using a Leica CM1850 UV cryostat and stored in −70°C, until analysis.

### Immunocytochemistry

After taking out from −70°C and drying for 15 minutes, slides were washed in PBS. The only time slides were fixed with 4% PFA 10 minutes in RT were for the Pax7/hLamininβ1/hLaminA/C/hSpectrin staining. Subsequently, slides were incubated for 1h at RT in blocking buffer containing 10% goat serum (Sigma, G9023) with 0.03 % triton X-100 in PBS and 1:50 Affini Pure Fab Fragment Donkey Anti-Mouse IgG (Jackson Immuno Research Laboratories, 715-007-003). Afterwards, slides were washed 5 minutes in PBS for 3 times and incubated with primary antibodies (Supplementary Table S4) in 10 % goat serum with 0.03% triton X-100 in PBS overnight at 4°C. The next day, slides were washed for 5 minutes in PBS for 3 times and incubated with secondary antibodies (Supplementary Table S5) in 10% goat serum with 0.03% triton X-100 in PBS for 1h in RT. Following washing for 5minutes in PBS for 3 times, slides were stained with DAPI 1:1000 solution in PBS and mounted in anti-fade fluorescence mounting medium (Abcam, ab104135). They were kept at 4°C prior to image acquisition using Cell DIVE Multiplex Slide Scanner (Leica Microsystems). For all transplant experiments, transverse sections with the most cells and myofibers of donor origin were identified and quantified.

### Hematoxylin and eosin (H&E) staining for slide sections

The H&E stain was done using an automatic staining machine Leica Multistainer ST5020 with Coverslipper CV5030.

## Data availability

All non-protected additional data are available from the authors upon request.

## Acknowledgements

We thank Penney Gilbert, Majid Ebrahimi and Ratima Suntornnond for helping fabrication of PDMS molds, William Weston for assistance in H&E staining, and Luke Gammon for help with Cell DIVE imaging. This research was primarily funded by the Barts Charity grant MGU0426 to YYL and JEM. AK was funded by the QMUL-Life Sciences Initiative PhD studentship and the Barts Charity. This work was in part supported by Royal Society grant RG130417, Newlife Charity grant SG/14-15/14, Action Duchenne grant AD1801Y, and Duchenne Parent Project grant 19.017 to YYL.

## Author Contributions

AK conducted most of the experimental work, analysed and interpreted the data. JM and JEM performed some of the experiments. JEM and YYL designed the experiments, analysed and interpreted the data, and supervised the research. OP and JC provided essential expertise, materials, and technical supports for the experiments. AK, JEM and YYL co-wrote the manuscript with inputs from all authors.

## Supplemental information

**Supplementary Table S1.** Quantification of transplantation experiments at 4 weeks.

| Experiment no. | Timepoint of TA collection | Condition | Number of cells/myofibers detected by human-specific antibodies |  |  |  |  |  |  |
| --- | --- | --- | --- | --- | --- | --- | --- | --- | --- |
|  |  |  | Undifferentiated cells |  | Differentiated human myofibers |  |  |  |  |
|  |  |  | hLaminA/C+ |  | hDys+ | hSpectrin+, hDys+, hLaminA/C+ | hSpectrin+, hDys+ | hSpectrin+, hLaminA/C+ | Total |
| 1 | 4 weeks | CORR-R3381X MPCs in Promocell | 2 |  | 0 | 2 | 2 | 0 | 4 |
|  |  |  | 4 |  | 0 | 5 | 0 | 0 | 5 |
|  |  |  | 1 |  | 0 | 2 | 0 | 0 | 2 |
|  |  |  | 1 |  | 0 | 0 | 1 | 0 | 1 |
|  |  |  | 10 |  | 1 | 16 | 10 | 0 | 27 |
|  |  |  | 1 |  | 0 | 2 | 0 | 0 | 2 |
|  |  | hCD133+ in Megacell | 6 |  | 1 | 26 | 17 | 0 | 44 |
|  |  |  | 18 |  | 9 | 45 | 49 | 0 | 103 |
|  |  |  | 9 |  | 0 | 1 | 1 | 0 | 2 |
|  |  |  | 36 |  | 47 | 156 | 74 | 0 | 277 |
|  |  |  | 4 |  | 11 | 22 | 20 | 0 | 53 |
|  |  |  | 16 |  | 9 | 85 | 28 | 0 | 122 |
| 2 | 4 weeks | CORR-R3381X MPCs in Promocell | 4 |  | 2 | 5 | 13 | 0 | 20 |
|  |  |  | 13 |  | 13 | 64 | 100 | 0 | 177 |
|  |  |  | 1 |  | 0 | 3 | 3 | 0 | 6 |
|  |  |  | 5 |  | 4 | 6 | 25 | 0 | 35 |
|  |  |  | 0 |  | 0 | 5 | 8 | 0 | 13 |
|  |  | CORR-R3381X MPCs in Megacell | 0 |  | 0 | 0 | 0 | 0 | 0 |
|  |  |  | 1 |  | 3 | 0 | 2 | 0 | 5 |
|  |  |  | 1 |  | 0 | 0 | 2 | 0 | 2 |
|  |  |  | 2 |  | 1 | 2 | 4 | 0 | 7 |
|  |  |  | 0 |  | 2 | 2 | 3 | 0 | 7 |
|  |  |  | 0 |  | 1 | 2 | 4 | 0 | 7 |
| 3 | 4 weeks | CORR-R3381X MPCs in Promocell | 7 |  | 1 | 5 | 13 | 1 | 20 |
|  |  |  | 10 |  | 1 | 16 | 22 | 0 | 39 |
|  |  |  | 12 |  | 2 | 29 | 66 | 2 | 99 |
|  |  |  | 4 |  | 3 | 23 | 88 | 1 | 115 |
|  |  | CORR-K295fs MPCs in Promocell | 6 |  | 1 | 5 | 19 | 0 | 25 |
|  |  |  | 0 |  | 0 | 20 | 40 | 0 | 60 |
|  |  |  | 3 |  | 0 | 3 | 7 | 0 | 10 |
|  |  |  | 2 |  | 0 | 3 | 10 | 0 | 13 |
|  |  |  | 2 |  | 0 | 5 | 19 | 0 | 24 |
| 4 | 4 weeks | CORR-K295fs MPCs in Promocell | 0 |  | 0 | 0 | 2 | 0 | 2 |
|  |  |  | 12 |  | 0 | 0 | 3 | 0 | 3 |
|  |  |  | 5 |  | 1 | 0 | 7 | 0 | 8 |
|  |  |  | 3 |  | 0 | 0 | 2 | 0 | 2 |
|  |  | CORR-K295fs MPCs in Megacell | 5 |  | 0 | 1 | 1 | 0 | 2 |
|  |  |  | 1 |  | 2 | 3 | 5 | 0 | 10 |
|  |  |  | 5 |  | 0 | 1 | 0 | 0 | 1 |
| 5 | 4 weeks | CORR-K295fs MPCs in Promocell | 1 |  | 0 | 0 | 0 | 0 | 0 |
|  |  |  | 4 |  | 0 | 0 | 0 | 0 | 0 |
|  |  |  | 3 |  | 0 | 1 | 0 | 0 | 1 |

**Supplementary Table S2.**
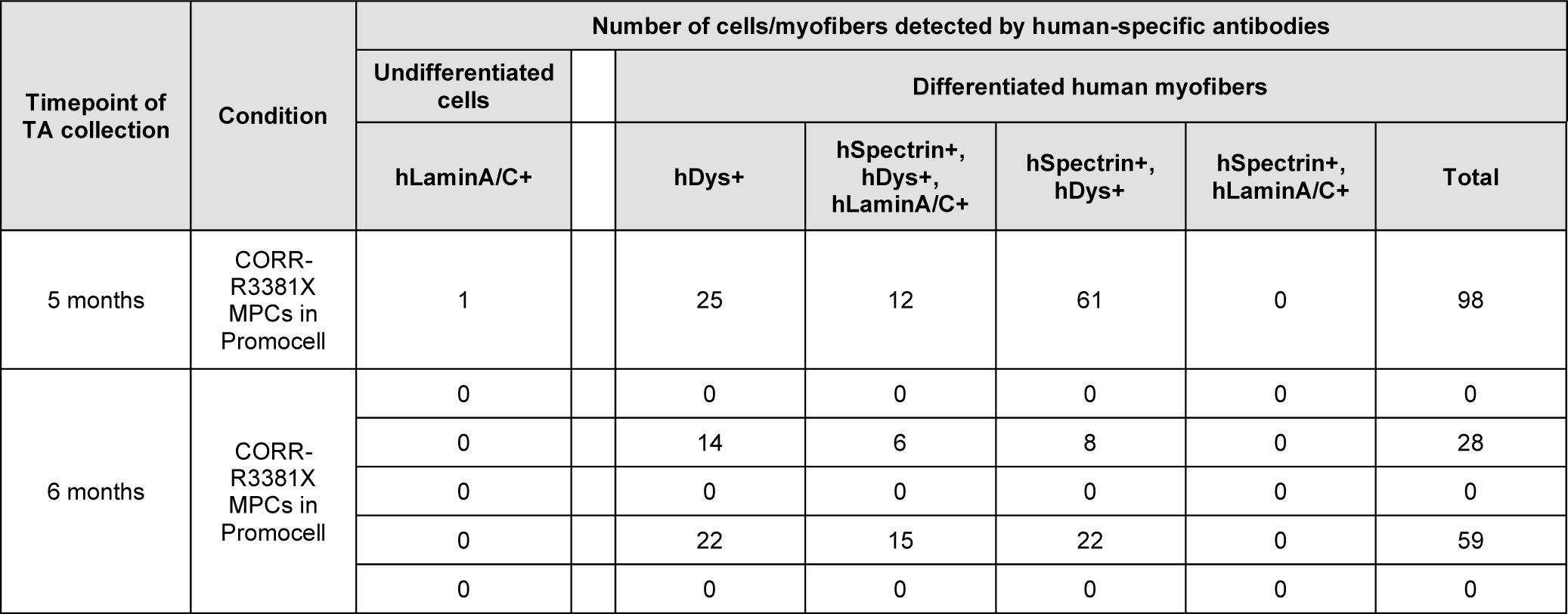
Quantification of transplantation experiments at 5 and 6 months.

| Timepoint of TA collection | Condition | Number of cells/myofibers detected by human-specific antibodies |  |  |  |  |  |  |
| --- | --- | --- | --- | --- | --- | --- | --- | --- |
|  |  | Undifferentiated cells |  | Differentiated human myofibers |  |  |  |  |
|  |  | hLaminA/C+ |  | hDys+ | hSpectrin+, hDys+, hLaminA/C+ | hSpectrin+, hDys+ | hSpectrin+, hLaminA/C+ | Total |
| 5 months | CORR-R3381X MPCs in Promocell | 1 |  | 25 | 12 | 61 | 0 | 98 |
| 6 months | CORR-R3381X MPCs in Promocell | 0 |  | 0 | 0 | 0 | 0 | 0 |
|  |  | 0 |  | 14 | 6 | 8 | 0 | 28 |
|  |  | 0 |  | 0 | 0 | 0 | 0 | 0 |
|  |  | 0 |  | 22 | 15 | 22 | 0 | 59 |
|  |  | 0 |  | 0 | 0 | 0 | 0 | 0 |

**Supplementary Table S3.** Reagents and cell culture medium recipes.

| # | Medium name | Details |
| --- | --- | --- |
| 1 | Promocell Skeletal Muscle Growth medium (C-23060) | 95 ml of basal medium (C-23060B), 5 ml of Supplement Mix (C-39365) and 105 µl 0.2% Penicillin/Streptomycin (Gibco, 15140122) |
| 2 | Megacell Skeletal Muscle Growth Medium | MegaCell Dulbecco's Modified Eagle's Medium (Sigma-Aldrich, M3942) supplemented with 10% fetal bovine serum (FBS, Gibco, 10270-098), 0.1mM β-Mercaptoethanol (Gibco, 31350-010), 1% NEAA (Gibco, 11140-035), 2µM Glutamine (GlutaMAX™ Supplement, Thermo Fisher, 35050061) and 5 ng/ml recombinant FGF-2 (PeproTech, 100-18B) |
| 3 | Fibrin/Matrigel hydrogel mixture | 4 mg/ml fibrinogen (40% vol; Sigma, F8630-1G), Matrigel (3.5 mg/ml; Fisher Scientific, 11543550) in DMEM/F12 media (40% vol; Invitrogen, 11320-033), thrombin (Sigma, T6634; 0.2 unit/mg fibrinogen) |

**Supplementary Table S4.** List of immunocytochemistry primary antibodies.

| # | Antibody | Isotype | Species | Dilution | Supplier | Cat no. |
| --- | --- | --- | --- | --- | --- | --- |
| 1 | Spectrin | IgG2b | Mouse | 1:50 | Leica Biosystems | SPEC1-CE |
| 2 | Lamin A/C | IgG2b | Mouse | 1:500 | Vector Labs | VP-L550 |
| 3 | Dystrophin (Mandys106; 2C6) | IgG2a | Mouse | 1:500 | Sigma-Aldrich | MABT827 |
| 4 | Pax7 | IgG1 | Mouse | 1:100 | DSHB | — |
| 5 | Laminin β1 | IgG1 | Mouse | 1:500 | Sigma-Aldrich | MAB1921P |
| 6 | CD31 (clone 390) | IgG2a | Rat | 1:100 | eBioscience | 14-0311-81 |
| 7 | SV2 (Synaptic vesicle glycoprotein 2A) | IgG1 | Mouse | 1:200 | DSHB | — |

**Supplementary Table S5.** List of immunocytochemistry secondary antibodies and conjugates.

| # | Antibody | Species | Dilution | Supplier | Cat no. |
| --- | --- | --- | --- | --- | --- |
| 1 | Anti-mouse IgG2a 488 | Goat | 1:1000 | Invitrogen | A-21131 |
| 2 | Anti-mouse IgG2b 594 | Goat | 1:1000 | Invitrogen | A-21145 |
| 3 | Anti-mouse IgG2b 546 | Goat | 1:1000 | Invitrogen | A-21143 |
| 4 | Anti-mouse IgG1 488 | Goat | 1:1000 | Invitrogen | A-21121 |
| 5 | Anti-rat IgG 546 | Goat | 1:500 | Invitrogen | A-11081 |
| 6 | α-Bungarotoxin<br>Conjugates, Alexa<br>Fluor 488 | — | 1:500 | Invitrogen | B13422 |

